# MaREA: Metabolic feature extraction, enrichment and visualization of RNAseq data

**DOI:** 10.1101/248724

**Authors:** Alex Graudenzi, Davide Maspero, Claudio Isella, Marzia Di Filippo, Giancarlo Mauri, Enzo Medico, Marco Antoniotti, Chiara Damiani

## Abstract

The characterization of the metabolic deregulations that distinguish cancer phenotypes, and which might be effectively targeted by *ad-hoc* strategies, is a key open challenge. To this end, we here introduce **MaREA** (Metabolic Reaction Enrichment Analysis), a computational pipeline that processes cross-sectional RNAseq data to identify the metabolic reactions that are significantly up-/ down-regulated in different sample subgroups. **MaREA** relies on the definition of a Reaction Activity Score, computed as a function of the expression level of genes encoding for reaction enzymes, which can also be used as an effective metrics to cluster samples into distinct metabolic subgroups. **MaREA** finally allows to visualize the results in a graphical form directly on metabolic maps. We apply **MaREA** to distinct cancer datasets and we show that it can produce useful information and new experimental hypotheses on metabolic deregulation of cancer cells, also allowing to stratify patients in metabolic clusters with significantly different survival expectancy.

## 1. Introduction

The heterogeneity of cancer genotypes and phenotypes hinders the identification of targets for effective treatments and is a major cause of tumor relapse. Therefore, it is common practice to statistically compare the gene expression of patient cohorts, based on clinical observations and/ or molecular features, in order to understand how the hallmarks of cancer can be (alternatively) achieved in terms of gene expression regulation. An hallmark in particular is of interest for cancer treatment: the *metmetabolic reprogramming* of cancer cells [1, 2].

Current research on cancer metabolism typically relies on genome-wide reconstructions of human metabolic networks [3], such as HMR [4] and Recon 2.2 [5]. These models include most metabolic reactions that may occur in a generic cell. Different strategies are used to extract relevant sub-models, e.g., the active metabolic networks in a given cell or tissue, in order to investigate its specific molecular features, possibly identifying relevant biomarkers and therapeutic targets. Current methodologies to this end employ transcriptome, proteome or even metabolome data (as reviewed in [6, 7, 8]). Such models are typically used in *Flux BalBalance Analysis* (FBA), a technique that exploits linear programming to compute the flux through each reaction under a *steady state assumption*, and which requires further constraints on incoming nutrients and outgoing products (i.e., exchange reactions) [9]. Yet, retrieving such information might be hard, as the simultaneous presence of metabolic measurements and transcriptomic data on same patient are rarely available in public databases, such as, e.g., the The Cancer Genome Atlas (TCGA) [10]

We here propose **MaREA (Metabolic Reaction Enrichment Analysis)** (Figure 1) as an alternative computational approach to investigate cancer metabolism, when data on metabolic measurements are not available, which explicitly focuses on *transcriptional regulation* of metabolic reactions.

**Figure 1:**
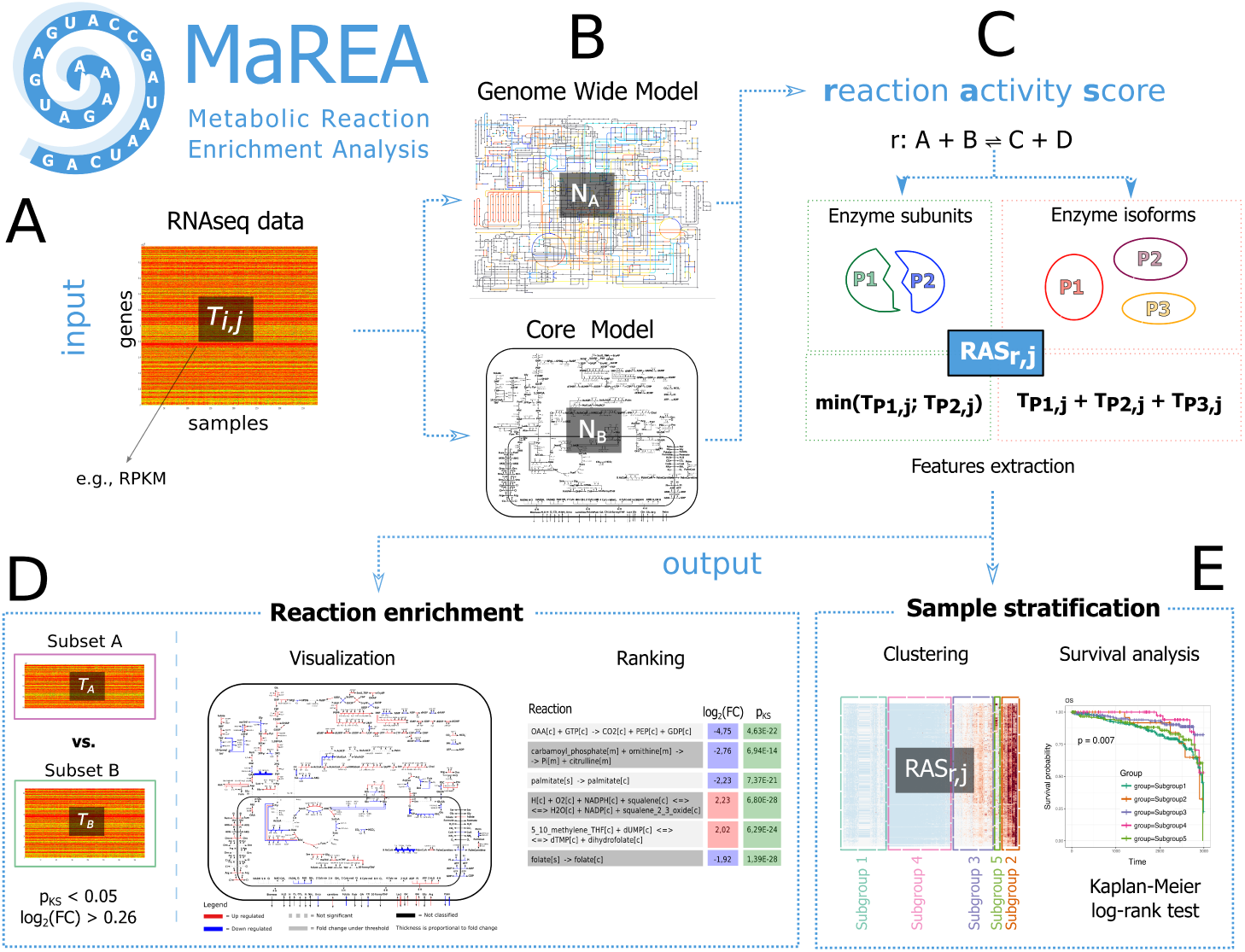
MaREA pipeline. **(A)** MaREA takes as input a *n* sample × m genes matrix *T* which includes the normalized read count of each gene from a given cross-sectional RNAseq dataset. **(B)** MaREA can use different input metabolic reaction networks, e.g., the genome-wide Model, or any subset of it. **(C)** A Reaction Activity Score (RAS) is defined for any reaction r in the input network and any sample, by distinguishing the case of reactions involving enzymes composed by different subunits - in this case the RAS is computed as the minimum of the transcript level of the genes encoding the subunits -, and that of reaction catalyzed by different enzyme isoforms - in this case the RAS is computed as the sum of the transcript level of the genes encoding the isoforms. **(D)** Given two distinct subsets *T_A_* and *T_B_* of the original dataset, the RASs of a given reaction in the two cases are compared and if the p-value of the Kolmogorov-Smirnov test is significant (< 0.05) and the log2 fold-change is larger than 0.263, that reaction will be enriched in the final graph as up‐ or down-regulated. Accordingly, a reaction ranking can be provided. **(E)** MaREA can stratify patients by employing the RAS as metrics on standard clustering methods, e.g., k-means. Survival analyses, such as log-rank test on Kaplan-Meier curves, can finally provide a prognostic validation of the clusters (notice that the image is intended for explanatory purpose only and does not reproduce any real case study).

**MaREA** processes *RNAseq data*, as one can retrieve from publicly available databases such as TCGA [10]. For each reaction of a given metabolic network, **MaREA** computes a *Reaction Activity Score* (RAS), which describes the extent of its activity in a given condition, as a function of the expression level of the genes encoding for the *subunits* and/ or the *isoforms* of the enzyme catalyzing such reaction. This score provides a more refined information than the mere list of genes associated to a reaction, without requiring to set any arbitrary threshold, nor to binarize data (gene present or absent), as required by other approaches [11]. Besides, **MaREA** does not perform FBA simulation, but only employs the RAS as a static representation of the metabolic behavior of a given sample, which can be then used to compare different sample sets (e.g., different patient cohorts, or control vs. tumor), identifying over (or under) expressed reactions.

Our approach markedly differs from that of *Gene Set Enrichment Analysis* (GSEA), which aims at characterizing the sets of up‐ or down-regulated genes in different phenotypes [12]. The typical outcome of GSEA analysis provides generic indications on the deregulated functions of a cell, or on specific functional behaviors when focusing on particular gene sets, derived, for instance, from the Reactome pathway database [13]. Nevertheless, the enriched sets are mere list of genes involved in comprehensive metabolic pathways (as, e.g., DNA replication), failing to provide details on which specific metabolic routes of a large pathway are favored in a given condition. In particular, metabolic functions can be alternatively achieved by metabolizing different nutrients and/ or by following different catabolic and anabolic routes, in a complex and largely undeciphered interplay. For this reason, simply knowing weather a certain function is *up or down-regulated* might not be sufficient to shed light on how such function might be achieved in distinct cancer phenotypes. In this regard, **MaREA** can provide a much finer resolution to the analysis and the enrichment of metabolic reactions in distinct experimental conditions.

More recent approaches aim at building sets of genes to be enriched according to the information included in genome-wide metabolic networks. Specifically, *metabolic reporter analyses* try to provide knowledge about variations in metabolite concentrations, starting from sets of genes classified according to the metabolite they associate with [14]. However, such methods do not provide information about which reactions are up‐ or down-regulated, and thus hinders the identification of putative targets for cancer treatment.

**MaREA** allows to overcome the limitations of current methods, providing a fine instrument for cancer metabolism investigation based on simple assumptions and easily-accessible transcriptomic data. Similar in spirit to the recently introduced PARADIGM approach (PAthway Recognition Algorithm using Data Integration on Genomic Models [15]), **MaREA** extracts a feature (the RAS) for each sample. However PARADIGM relies on the integrating of data from multiple sources and requires curated pathway interactions among genes, hence both the input data and final objective are significantly different.

Importantly, the features extracted by **MaREA** can be used to stratify samples in an unsupervised manner (*Metabolic Feature Extraction*). Such stratification might provide relevant prognostic indications, as shown in the case studies in Section 2. Distinct approaches make use, for instance, of the information on enriched pathways [16] or of that on mutational profiles [17, 18] to classify cancer samples and subtypes.

Summarizing, **MaREA** can be used to: *i*) rank the reactions according to the variation in their activity observed between different phenotypes and/ or experimental conditions; *ii*) enrich the map of human metabolic routes with the variation observed in the RAS of each reaction, providing a clear visualization of how deregulated paths are interconnected; *iii*) efficiently stratify samples according to their metabolic activity, hence providing a new (unsupervised) clustering tool, with testable clinical relevance, which can be assessed, e.g., via standard survival analyses.

As a proof of principle, we applied **MaREA** to data publicly available in the TCGA database [10]. We computed the RASs by taking into account either all the Gene-Protein Rules (GPRs) included in the genome wide-model Recon 2.2 [5] or a manually curated subset of it, corresponding to the model of central carbon metabolism previously used in [19, 20]. We identified and visualized the enriched metabolic reactions between normal and cancer biopsies obtained from breast cancer patients (TCGA-BRCA dataset [21]). Furthermore, we used the RASs to stratify breast cancer patients in distinct metabolic clusters, and performed standard survival analysis, which highlighted statistically significant prognostic predictions.

## 2. Materials and Methods

*Input*. **MaREA** takes as input any RNAseq dataset in the form of a *n* × *m* matrix *T*, where *n* is the number of genes and *m* is the number of samples of the considered cohort (see Figure 1-A). Each element *T*_*i*,*j*_, *i* = 1,…, *n*, *j* = 1,…, *m* corresponds to the *normalized read count* of gene *i* in sample *j* such as, for instance, the RPKM (Reads per Kilobase per Million mapped reads).

**MaREA** then filters *T* according to a specific input reaction network *N*, e.g., the genome-wide metabolic network [5] or any possible subset of it (see Section 2). In particular, we define the set of reactions as *R* = {*r* ∈ *N*}. Therefore, *T* is filtered by retaining only the rows corresponding to genes that are associated to enzymes involved in the reactions included in *R* (see Figure 1-B).

Gene-protein rules are logical formulas that describe how gene products concur to catalyze a given reaction. Such formulas include *AND* and *OR* logical operators. AND rules are employed when distinct genes encode different *subunits* of the same enzyme, i.e., *all* the subunits are *necessary* for the reaction to occur. OR rules describe the scenario in which distinct genes encode *isoforms* of the same enzyme, i.e., either isoform is *sufficient* to catalyze the reaction.

For example the succinate-Coenzyme A ligase enzyme is formed by the subunits alpha (gene SUCLG1) and beta gene (SUCLG2) and catalyzes the reaction *Pi* + *succinyl-CoA* + *GDP* ↔ *CoA* + *succinate* + *GTP*. The gene-enzyme rule for this reaction is therefore: SUCLG1 *AND* SUCLG2. Conversely, ACACA and ACACB are respectively fully functional enzyme for the reaction *acetyl-coenzyme A ligase carboxylase*, thus the rule is ACACA *OR* ACACB. Such logical operators can of course be combined to depict multi-protein catalytic complexes or more complex situations involving both subunits and isoforms. For instance, ribonucleotide reductase is formed by two subunits: the catalytic (M1) and the regulatory one. The latter exists in two isoforms (M2 and M2B). The rule for this enzyme will therefore be RRM1 *AND* (RRM2 *OR* RRM2B).

### Reaction Activity Score (RAS)

To avoid the definition of arbitrary thresholds on the transcript level, we do not resolve the logical expressions in a Boolean fashion, but we define a *Reaction Activity Score* (RAS), for each sample *j* = 1,…,*m*, and each reaction *r* ∈ *R* (see Figure 1-C). In order to compute the RAS we distinguish:

- Reactions with AND operator (i.e., enzyme subunits).

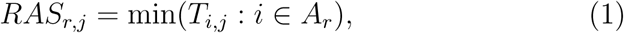

where *A*_*r*_ is the set of genes that encode the subunits of the enzyme catalyzing reaction *r*.
- Reactions with OR operator (i.e., enzyme isoforms).

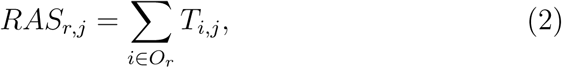

where *O*_*r*_ is the set of genes that encode isoforms of the enzyme that catalyzes reaction *r*.

In case of composite reactions, we respect the standard precedence of the two operators. The final output is therefore a |*R*| × *m* matrix *M*, where each element *M*_*r*,*j*_ is the RAS computed for reaction *r* in sample *j*.

The intuition underneath the introduction of the RAS is that enzyme isoforms contribute *additively* to the overall activity of a given reaction, whereas enzyme subunits *limit its activity*, by requiring all the components to be present for the reaction to occur.

Clearly, we are here adopting a deeply simplified approach to reaction network modeling, by neglecting, for instance, the great heterogeneity of reaction kinetic constants and protein binding affinities, of translation rates, and any possible post-transcriptional regulation effect that might occur within a cell. In this regard, an optimal choice would be to weigh all the reactions according to such quantities, yet direct measurements or robust estimates are very rarely available, especially for genome-wide models. Therefore, on first approximation, we here assume that all enzyme isoforms and subunits contribute uniformly to the reaction activity of a given reaction, as we expect that this choice does not affect the up-/ down-regulation interplay observed at the network level.

### Reaction enrenrichment: visualization and ranking

One important outcome of the RAS introduction is the possibility of identifying and visualizing in an explicit way the metabolic routes that are up‐ or down-regulated in different sample sets and/ or experimental conditions (see Figure 1-D).

Given two distinct RNAseq datasets, or two partitions of the same dataset, *T*_*A*_ and *T*_*B*_, and an input metabolic reaction network *N*, we first compute the RAS matrices *M*_*A*_ and *M*_*B*_. For each reaction *r* ∈ *N* we then perform a non-parametric two-sample *Kolmogorov-Smirnov* (KS) test with a standard p-value threshold of 0.05, to verify whether the distributions of RASs over the samples in the two sets are significantly different.

In that case, we compute the *log*_2_ fold-change of the average RAS*_r_* in the two groups. Because KS-test considers as significantly different distributions with the same mean, but different standard deviation, we consider as relevant only *log*_2_ fold-change larger than 0.263 (i.e., corresponding to a 20% variation of the average RAS). In line with the philosophy of GSEA[12], we use a relaxed threshold for the fold-change, because even an increase of 20% in genes encoding members of a metabolic pathway may dramatically alter the flux through the pathway. The significance threshold on the p-value of the Kolmogorov-Smirnov test, on the other hand, ensures that expression distributions of the two groups are indeed different and, therefore, that a 20% difference in the average expression level does not occur by chance.

**MaREA** then uses the significant RAS fold-changes to: *i*) determine a ranking of the most relevant up‐ and down-regulated reactions in the two sets, *ii* map such quantities over the input metabolic network *N*, by respectively coloring in red/ blue the up-/ down-regulated reactions, and by setting the edge thickness as proportional to the RAS fold-change. The reactions that will either display non-significant p-value or a RAS fold-change below the threshold will not be included in the ranking and will be marked in gray color on the metabolic network.

### RAS-basedsample stratification

Another major advantage of our approach is that it is possible to employ the RAS as an efficient metric to identify sample subgroups (or *clusters*) that share similar metabolic properties (see Figure 1-E). In particular, we here implemented a *k-mk-meanseans* clustering [22] which use the RASs of all reactions *r* ∈ *R* to identify sample clusters with distinct metabolic behaviours. Clusters can be compared by means of the reaction enrichment procedure described above, by ranking the significantly different reactions in the distinct clusters and visualizing the RAS fold-changes on the input network. Above all, clusters can also be tested via standard *survival analysis*, such as the *log-rank* test on *Kaplan-Meier* curves (when data is available), hence providing an important orthogonal validation of the clustering results with clinical relevance. In Section 2 we show how clustering on RASs indeed can produce significant prognostic predictions.

### Metabolic Network

To compute the RAS of cancer samples in a given TCGA dataset at the genome-wide scale, we used the GPRs included in the most up-to-date genome-wide network of human metabolism:*Recon* 2.2 [5]. In particular, to visualize **MaREA** results at the genome-wide level, we modified the graphical attributes of the model map in *xml*format obtained from the *Virtual Metabolic Human* (VMH) (https://vmh.uni.lu/ which is readable by the tool *Cell Designer* [23].

To focus, instead, on central carbon metabolism, we used the metabolic core model(*HMRcore*) introduced in [20]. For the sake of completeness, we included in the model mitochondrial palmitate degradation and gluconeogenesis. As the original version of the model does not include information on GPR_s_, such rules have been extracted and manually curated from Recon 2.2 [5] and included in the HMRcore model. In particular, we verified the correctness of gene-protein rules taking into account the information retrieved from the *Human Protein Atlas* [24] for the protein tissue location, from *UniProtKB* [25] for the enzyme complex composition, and from *KEGG* [26] to check gene - enzyme association. Notably, we corrected for some inconsistencies within Recon 2.2, with particular regard to cytosolic reactions associated with mitochondrial isoforms or the other way around. The final version of the HMRcore model includes 231 reactions with rules and 390 metabolic genes that are associate to them. Genes are identified with the *HGNC ID* provided by the *HUGO Gene Nomenclature Committee* [27]. The SBML of the model is provided in Supplementary file S1.

It should be noted that not every reaction in the metabolic models is associated with a gene-enzyme rule: for instance some reactions have been included to fill the gaps in steady state computations, but we lack knowledge on the associate genes. In detail, 4742 (231) reactions over 7785 (272) are associated with a gene-enzyme rule, in the genome-wide (core) model and thus a RAS can be computed for them.

### Dataset

We applied the **MaREA** pipeline to the breast cancer dataset (TCGA-BRCA) published in [21], which also includes healthy/ control samples. We downloaded the dataset via the *cBioPortal* [28] (case study id: brca_tcga_pub2015). This dataset includes the expression profile (RNA Seq V2 RSEM) of biopsies taken from 817 patients. For 105 of them, the expression profile of the normal tissue is also included. Because the TCGA-BRCA dataset identifies genes with Entrez IDs, we automatically converted them into *HGNC* IDs.

We found a correspondence for 1654 (379) genes over the 1673 (390) included in the genome-wide metabolic model. Although we neglected missing genes in the computation of the RAS_s_, we were still able to compute a RAS for each reaction associated with a GPR.

## 3. Results

### 3.1. Breast Cancer vs.Normal

In order to evaluate the goodness of **MaREA** results, we first applied it to a largely characterized case-study: the comparison of cancer and normal metabolism.

#### Reaction enrichment

The reactions that have been identified by **MaREA** as significantly up or down regulated in cancer - with at least a 20% increase/decrease - and the magnitude of the deregulation are mapped on the central carbon metabolic network (HMRcore) in Figure 2, as well as on the genome-wide metabolic network in Supplementary Figure S1. It can be easily observed (Figure 2) that the pathway of glycolysis is over expressed in cancer. Extensive utilization of glucose is indeed a well established trait of breast and of cancer cells in general [29]. Cancer cells need glucose to feed the metabolic requirement of enhanced proliferation, with particular regard to: 1) *de novo* synthesis of nucleotides for genome replication; 2) synthesis of amino acids for protein synthesis 3) synthesis of fatty acids to support the expansion of cellular membranes; 4) ATP generation for energetic requirements. Accordingly, **MaREA** returned as largely up regulated in cancer: 1) synthesis of nucleotides from Phosphoribosyl-pyrophosphate (PRPP in Figure 2); 2) metabolism of the non-essential aminoacids serine (Ser), glycine (Gly), alanine (Ala), asparagine (Asn), aspartate (Asp), arginine (Arg) and proline (Pro); 3) synthesis of cholestrol (Chol) from citrate (Cit).

**Figure 2:**
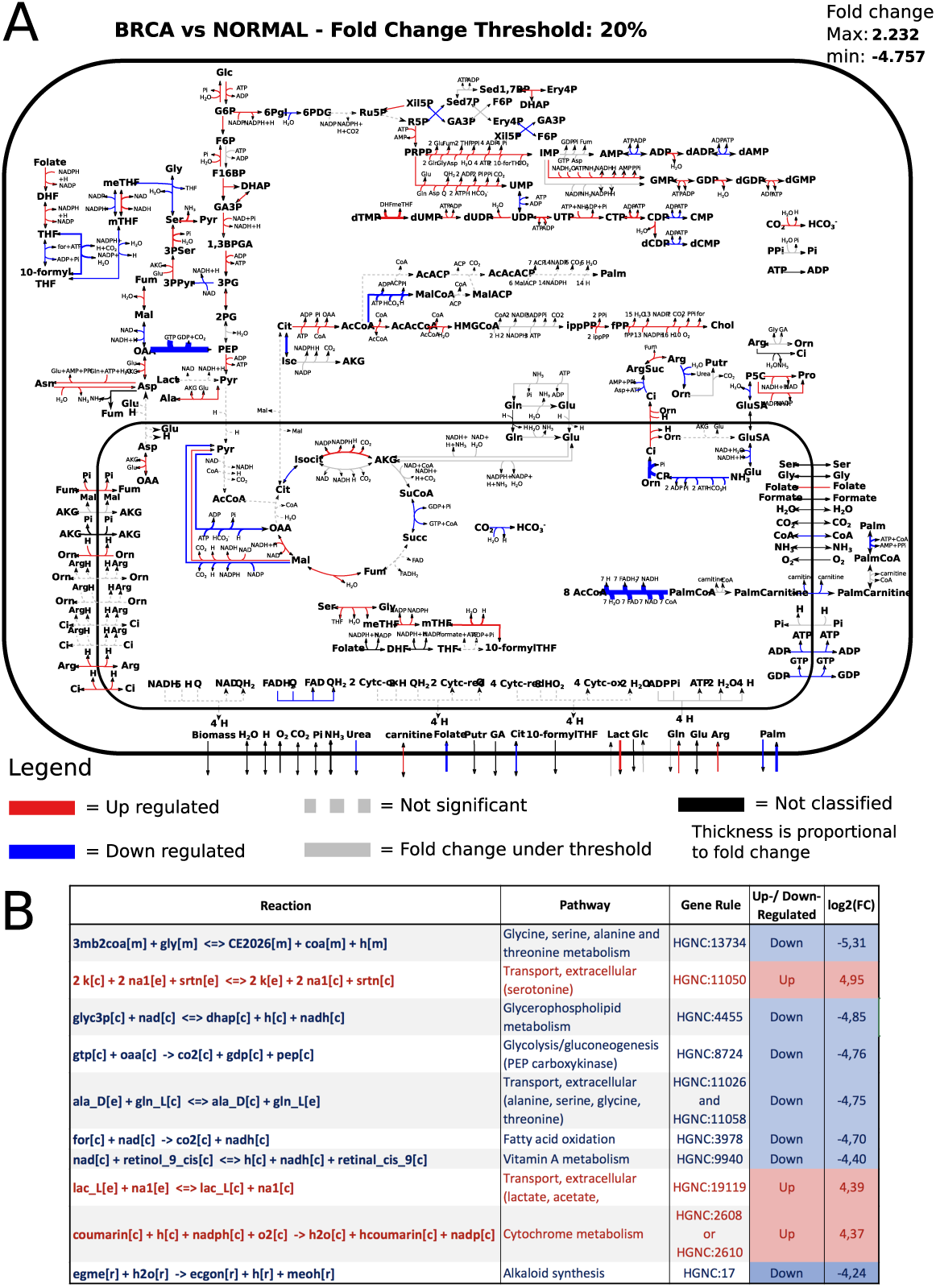
Breast cancer vs. normal samples. **(A)** HMRcore map enriched by MaREA: Reactions up-regulated in breast cancer sample set are marked in red, reactions up-regulated in normal sample set are marked in blue. A list of the abbreviations used in the map is provided in Supplementary Text S1. Thickness of the edges is proportional to the fold-change. Non-Classified reactions, i.e., reactions without information about the corresponding gene-enzyme rule, are marked in black. Dashed gray arrows refer to non-significant deregulations according to the Kolmogorov-Smirnov test. Solid gray arrows refer to reactions with a log2 fold-change below 0.263. **(B)** A reaction ranking is provided, by listing the 10 reactions with largest log2 fold-change of the RAS (absolute value) in the two conditions. The reaction formula, the corresponding pathway, the gene rule, the up-/ down-regulation flag and log2 fold-change are shown in the table.

As long as ATP production is concerned, the interpretation of the situation portrayed by **MaREA** is, as expected, less straightforward. Cancer cells are believed to rely more on fermentation of glucose to lactate rather than on oxidation of glucose in the mitochondria (inner box of the map in Figure 2), despite the presence of oxygen: a phenomenon well-known as the Warburg Effect [2, 1]. However, in contrast to Warburg'’s original hypothesis that damaged mitochondria are at the root of this phenomenon, the ability of mitochondria to carry out oxidative phosphorylation is not defective in most tumors [1]. In line with these studies, if on the one hand lactate secretion seems to be up regulated in cancer (reaction crossing the external box in Figure 2), on the other hand the respiratory chain (represented by the 5 reactions at the bottom of the mitochondrial box, which are catalyzed by protein Complexes I-V) is not significantly down-regulated in this breast cancer dataset, exception made for Complex II.

It is worth noticing that oxidation of NADH (Complex I and III-V) may occur independently from oxidation of FADH2 (Complex II and 3-V) in the respiratory chain, provided that NADH is not produced by a cyclic activity of the tricarboxylic acid (TCA) cycle. Remarkably, results in Figure 2 suggest that the working-mode of the TCA cycle may be abnormal in breast cancer. In particular, up regulation of NADPH-dependent isocitrate dehydrogenase, which catalyzes the reductive carboxylation of ‐ketoglutarate (AKG) to isocitrate, may be linked with the mutations often reported for this enzyme in breast and other cancer types [30]. It has been suggested that this enzyme may support reductive glutamine metabolism in cancer and a branched TCA cycle flux mode [31]. Enhanced utilization of glutamine is indeed another hallmark of cancer cells [1, 2]. Accordingly, intake of glutamine is reported to be up regulated by **MAREA**.

The agreement of the results in Figure 2 with the obvious traits of cancer metabolism supports the reliability of our approach, which might shed light on less established traits. Deregulations of breast cancer metabolism identified by the approach, which may be worth of note are, among others: 1) deregulation of beta-oxidation of palmitate; 2) upregulation of folate metabolism; 3) deregulation of Phosphoenolpyruvate carboxykinase, which converts oxaloacetate (OAA) into phosphoenolpyruvate (PEP).

#### Reaction ranking

After filtering out the reactions whose activity does not differ between cancer and normal samples, **MaREA** allows to rank the remaining reactions according to the extent of their up‐ or down-regulation. Supplementary Table S1 report, for each reaction included in the genome-wide model: the log fold change, the reaction formula, the pathway in which the reaction is involved and a description of its role. As genome-wide models include several reactions that are associated with the very same GPRs, typically involving transporters/enzymes with low substrate specificity, in Figure 2 we report the top 10 deregulated reactions with different GPRs.

These reactions include 3 up regulated reactions and 7 down-regulated ones. Most of theme are associated with a single gene, including, consistently with the results obtained for the HMRcore model: fatty acids oxidation and PEP carboxykinase, which are significantly down regulated; lactate (or substrates pertaining to the same family) transport, which is up-regulated in cancer. Single-gene top deregulated reactions not included in the HMRcore model relate to deregulated vitamine A, glycine and alkaloid metabolism and to upregulated transport of serotonine. Notably, in accordance with this results, it has been reported [32] that serotonine promotes tumor growth and survival in breast cancer, and that vitamin A [33] plays a role in cancer treatment and prevention.

Two top deregulated reactions are associated with a pair of genes linked by an OR and AND respectively: 1) down regulated antiporter of the aminoacids alanine, serine, glycine and threonine with glutamine; 2) up regulated Cytochrome P450 2A6, which is involved in the metabolism of many xenobiotics.

The down-regulation observed for the former reaction is in line with recent studies that have linked the resistance of specific cancer cell lines to amino acid analogs anticancer drugs to a decreased expression of the corresponding transporter [34, 35].

The up-regulation identified for the latter reaction (P450 2A enzyme) is worth of note, as P450 enzymes may be involved in carcinogens activation in breast cancer. Environmental carcinogens have been identified in the etiology of breast cancer. For example, CYP2A6 protein detected in the breast can activate nitrosamines and food mutagens to their ultimate carcinogens and thus could play a role in the initiation of breast cancer [36]. Moreover, this enzyme can metabolize clinically important drugs, such as the tamoxifen [37], which represents the most widely used hormonal therapy for breast cancer, and the coumarin [38, 39], whose metabolisms have been proven to produce some metabolites having estrogenic and cytotoxic activities.

#### Comparison with GSEA results

The GSEA and **MaREA** approaches are not directly comparable, as they present several differences in goals, input data, parameters, variables and outputs. For instance, MAREA computes an individual activity score for each sample, whereas GSEA only considers expression fold-changes between pairs of experimental conditions. In order to provide an overview of how the information produced by the two complementary approaches may differ, without the ambition of claiming which approach should be preferred, we disregarded the addition multiple test correction (FDR) used by GSEA, and we considered the gene-sets that pass the nominal p-value test, with the same threshold used in MAREA standard settings (i.e., p=0.05). We did not set any threshold on the minimum size of gene-sets.

To run GSEA we used two kind of gene sets: 1) curated gene sets based on REACTOME, as directly provided by the GSEA tool; 2) gene-sets reconstructed by us, which correspond to the genes associated to each reaction in the genome-wide model and which are provided in *gmt* format (GSEA compliant) in Supplementary File S2. The former kind of gene-sets represent a typical application of GSEA to gene-sets involved in broad metabolic functions; whereas the second type is directly comparable to the sets used to compute the RAS by our approach. It should be mentioned that the second type includes many single gene-sets (size 1), because many reactions are catalyzed by enzymes associated to a single gene.

The application of GSEA to Reactome gene-sets returned 144 gene sets significantly enriched in cancer, and 60 gene-sets significantly enriched in normal, at nominal p-value < 0.05. When ranking the obtained gene-sets according to the returned Enrichemnt Score (ES), we observed that, as expected, the first 10 gene sets enriched in cancer refer to generic metabolic functions (in particular: cell cycle and mitosis, asparagine glycosylation, DNA replication and chromosome maintenance, HIV infection and kinesins). The results highlight how **MaREA** should be used as a complement to GSEA analysis, in order to provide a more fine-grained analysis of metabolic deregulations.

On the other hand, the application of GSEA to the more fine-grained datasets, based on RECON 2.2 reactions, returned a number of reactions significantly deregulated much lower than that returned by **MaREA**. **MaREA** returned 3339 reactions as significantly up‐ or down-regulated by at least 20% (p-value < 0.05); whereas GSEA returned 105 gene sets significantly enriched in cancer, and 110 gene-sets significantly enriched in normal, at nominal p-value < 0.05. This discrepancy is mainly due to the presence of gene-sets including a single-gene, which are reasonably penalized by the GSEA approach. By way of example, the single-gene sets associated to serotonin and vitamin A metabolism, which were ranked in the top 10 deregulated reactions by **MaREA**, and which might play a role in cancer according to literature as described above, do not pass the nominal p-value test in GSEA. It is worth noticing that also the top-ranked reactions in **MaREA** that involve gene in *OR* (Cytochrome metabolism) or in *AND* (extracellular transport of alanine, serine, glycine and threonine) do not pass the significance test in GSEA.

Taken together, these results indicate that **MaREA** provides a more complete and refined portray of metabolic deregulations. Moreover, as opposed to GSEA, **MaREA** computes an independent score for each sample (the RAS), which can be used to cluster samples in an unsupervised fashion. We illustrate such application of **MaREA** in the next section.

### 3.2. Metabolic Subgroups of Breast Cancer

In order to stratify samples by employing the RAS as clustering metrics, we performed *k-means* clustering with different input *k* ∈ {1, 2,…, 9} on the normalized RASs^1^ of the reactions included in the HMRcore model. We performed *n* = 100 bootstrap iterations, with random centroid assignments, selecting as optimal the clustering run displaying the maximum inter-cluster distance. We then tested the resulting sample clusters against the survival probability (as retrieved from clinical data in the original dataset [21]), via log-rank test on Kaplan-Meier curves.

Interestingly, we found statistically significant *overall survival* curves (*p* < 0.05) for *k* = 2, *k* = 3, and *k* = 9 (significant results with respect to disease-free and progression-free survival curves were observed as well, results not shown here).

In Figure 3 we show the most interesting result, obtained with *k* = 3 (Subgroup 1: 94/817 samples, Subgroup 2: 544/817 samples, Subgroup 3: 179/817 samples). One can see that the curves of the three clusters never overlap, leading to a highly significant log-rank test (*p* = 0.013). This result indicates that the up-/ down-regulation patterns, as encoded by the RAS values, might indeed be used to split samples in metabolic groups with significantly different prognosis. It is also worth noticing that, by looking at the composition of the three subgroups as computed on the 481 samples with respect to the well-established PAM 50 classification [40] (on 817 of the whole dataset), the subgroup with the worst prognosis (Subgroup 3) is largely constituted (i.e., ~ 70%) by samples belonging to the Basal-like group, which are completely absent from Subgroup 1 (best prognosis) and present in very small percentage in Subgroup 2. Subgroup 1 and 2 are, instead, composed by a more complex mixture of samples from Luminal A, Luminal B and Her2 subtypes in different proportions.

**Figure 3:**
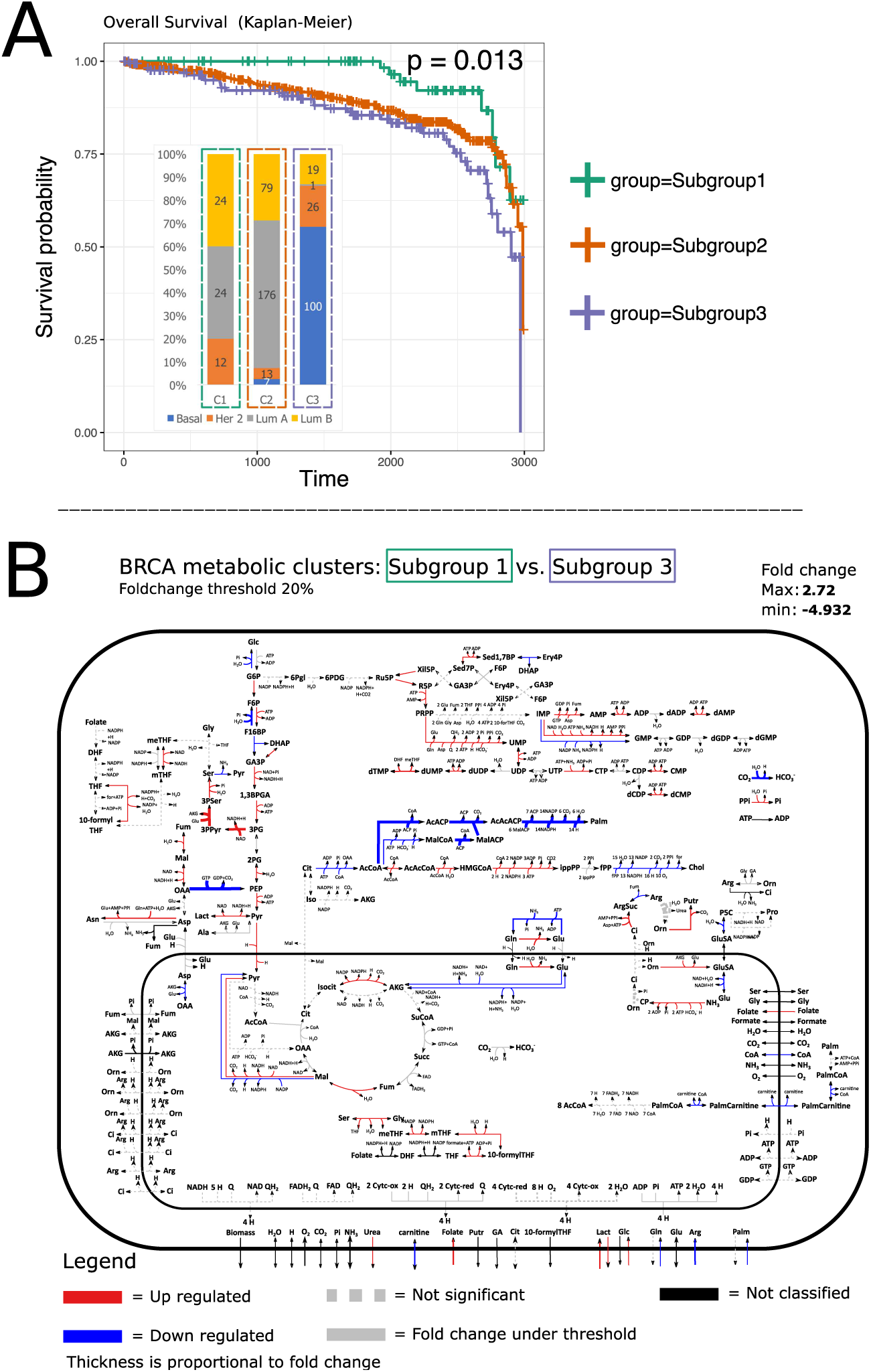
Breast cancer metabolic clusters. **(A)** The Kaplan-Meier curves (time unit = days) and the p-value of the log-rank test with respect to the three metabolic clusters of TCGA-BRCA samples identified by MaREA are shown (Subgroups 1, 2 and 3). In the histograms the composition of the three clusters with respect to the standard PAM 50 classification (Basal-like, Her 2 positive, Luminal A and Luminal B) is provided. The numbers on the bars indicate the absolute value of samples in each subgroup. **(B)** Enriched map of HMRcore with respect to metabolic clusters 1 and 3. A list of the abbreviations used in the map is provided in Supplementary Text S1. Red arrows refer to reactions upregulated in Subgroup 3; whereas blue arrows refer to reactions upregulated in Subgroup 1. Black arrows refer to Not Classified reactions, i.e., reactions without information about the corresponding gene-enzyme rule. Dashed gray arrows refer to non significant deregulations according Kolmogorov-Smirnov test. Solid gray arrows refer to reactions with a log2 fold change below 0.263.

This result would first suggest that there exist a detectable metabolic *signature* of Basal-like cancer samples, and this is, to the best of our knowledge, a novel result, worth of further investigations. Besides, the mixed composition of Subgroups 1 and 2 may suggest that the differences observed at the metabolic level indeed translate into distinct survival probabilities, which standard classification may fail to capture.

More in detail, by looking at the reactions significantly up-/ down-regulated with respect to the case Subgroup 1 (best prognosis) vs. Subgroup 3 (worst prognosis) portrayed in Figure 3 for the core model - and in Supplementary Figure S2 and Table S2 for the genome-wide model - one can see that many reactions that are enriched in cancer against normal are also enriched in worst against best prognosis, including glycolysis, nucleotide synthesis and serine metabolism. Remarkably some metabolic pathways that are not significantly deregulated in cancer are significantly up regulated in the worse prognosis subgroup, with particular regard to palmitate biosynthesis.

## 4. Conclusions

We have here introduced **MaREA**, a computational pipeline that processes trascriptome profiles to produce usable information on metabolic reaction activities in different sample subgroups or experimental conditions. This is made possible thanks to the introduction of the Reaction Activity Score, which is computed as a function of the expression level of genes involved in reaction catalysis.

Thus, the resulting enriched reactions can provide a metabolic-centered explanation of the different phenotypic/ functional properties observed in distinct sample subgroups or cancer subtypes. The interpretation of the results is then favored thanks to the effective visualization of up-/ down-regulated reactions directly on the metabolic networks.

The case studies on TCGA cancer datasets proved that **MaREA** can reproduce known properties and traits of metabolic networks in different scenarios, for instance, by identifying the key metabolic paths that distinguish normal from tumor samples, but it can also provide cues to formulate and test new experimental hypotheses (e.g., relevance of serotonine metabolism in breast cancer, see Figure 2).

Finally, **MaREA** allows to effectively identify metabolic clusters of samples, which display significantly different survival expectancy, as retrieved from clinical data. By relying on core models, **MaREA** allows to reduce the input dataset dimensionality, from thousand genes to the (much) fewer genes involved in the regulation of a given metabolic network. Therefore, **MaREA** might represent a powerful tool to link the metabolic behaviour to particular prognoses or to tumor aggressiveness, without employing any further genomic or molecular information.

It goes without saying that **MaREA** does not provide information on metabolic fluxes. For a deeper understanding of cancer metabolism, **MaREA** results should thus be complemented with metabolic measurements and flux simulations.

We remark that the final quality of **MaREA**'’s results deeply depends on the correctness of the gene-protein rules. We provided a curated version of the core model, but we automatically derived such rules from Recon 2.2 for the genome-wide analysis, thus our results might be improved via a more refined manual curation.

Even though we have here showed an application of **MaREA** to cancer patients, we remark that the approach can be used to compare any pair of conditions described by RNA-seq data samples (as, e.g., wild type vs. mutant organism). In the next future, we plan to release an user-friendly tool, which may be integrated within the Galaxy platform [41], to leverage the application of **MaREA** to any case study.

## Acknowledgments

This work is supported with FOE funds from the Italian Ministry of Education, Universities and Research (MIUR, http://www.http://www.istruzione.it/istruzione.it/) to SYSBIO - within the Italian Roadmap for ESFRI Research Infrastructures; by grants from AIRC (investigator grant IG 16819 and 9970-2010 Special Program Molecular Clinical Oncology 5x1000) to EM; and by funds from the FLAG-ERA grant ITFoC to GM. The funding sources had no role in study design, data collection and analysis, decision to publish, or preparation of the manuscript.

## Footnotes

1 To avoid possible biases due the differences in RAS range and distribution across reactions, we here normalize the RAS value of each sample by dividing by the maximum RAS for that reaction.

